# Non-coding Somatic Mutations Converge on the PAX8 Pathway in Epithelial Ovarian Cancer

**DOI:** 10.1101/537886

**Authors:** Rosario I. Corona, Ji-Heui Seo, Xianzhi Lin, Dennis J. Hazelett, Jessica Reddy, Forough Abassi, Yvonne G. Lin, Paulette Y. Mhawech-Fauceglia, Jenny Lester, Sohrab P. Shah, David G. Huntsman, Alexander Gusev, Beth Y. Karlan, Benjamin P. Berman, Matthew L. Freedman, Simon A. Gayther, Kate Lawrenson

**Affiliations:** Women’s Cancer Program at the Cedars-Sinai Samuel Oschin Comprehensive Cancer Center, Los Angeles, CA, USA; Center for Bioinformatics and Functional Genomics, Cedars-Sinai Medical Center, Los Angeles, CA, USA; Department of Medical Oncology, Dana-Farber Cancer Institute, Boston, MA, USA; Center for Functional Cancer Epigenetics, Dana-Farber Cancer Institute, Boston, MA, USA; Department of Obstetrics and Gynecology, Keck School of Medicine, University of Southern California, Los Angeles, CA; Aurora Diagnostics, Austin, TX, USA; Department of Computer Science, University of British Columbia, Vancouver, BC, Canada; Department of Molecular Oncology, BC Cancer Agency, Vancouver, BC, Canada; Department of Pathology and Laboratory Medicine, University of British Columbia, Vancouver, BC, Canada; Department of Gynecology and Obstetrics, University of British Columbia, Vancouver, BC, Canada; McGraw/Patterson Center for Population Sciences, Dana-Farber Cancer Institute, Boston, MA, USA; The Eli and Edythe L. Broad Institute, Cambridge, MA, USA

## Abstract

Transcriptional regulation is highly disease and cell-type specific. We performed H3K27ac chromatin immunoprecipitation and transcriptomic sequencing in primary tumors for the four different subtypes of invasive epithelial ovarian cancer (OC). Histotype-specific regulatory elements (REs) were enriched in enhancers (P<0.001). *In silico* prediction of putative target genes for histotype-specific REs identified genes (*WFDC2*, P=5.5×10^-5^) and pathways (PI3K-Akt signaling, P<0.002) known to be involved in OC development. Some genes (e.g. *PAX8* and *CA125*) are associated with super-enhancers (SEs) in all OCs, while others are histotype-specific, including *PPP1R3B* which is associated with SEs specific to clear cell OC. Integrated analysis of active chromatin landscapes with somatic single nucleotide variants (SNVs) from whole genome sequencing (WGS) of 232 primary OCs identified frequently mutated REs, including the *KLF6* promoter (P=8.2×10^-8^) and a putative enhancer at chromosome 6p22.1 (P<0.05). In high-grade serous OCs, somatic SNVs clustered in binding sites for the PAX8 binding partner TEAD4 (P=6×10^-11^), while the collection of *cis* regulatory elements associated with *PAX8* was the most frequently mutated set of enhancers in OC (P=0.003). Functional analyses supported our findings: Knockdown of *PPP1R3B* in clear cell OC cells significantly reduced intracellular glycogen content, a signature feature of this histotype; and stable knockout of a 635 bp region in the 6p22.1 enhancer induced downregulation of two predicted target genes, *ZSCAN16* and *ZSCAN12* (P=6.6 x 10^-4^ and P=0.02). In summary, we have characterized histotype-specific epigenomic and transcriptomic landscapes in OC and defined likely functional REs based on somatic mutation analysis of ovarian tumors.

## INTRODUCTION

Epithelial ovarian cancer (OC) is a heterogeneous disease, comprising several different histological subtypes that differ in their underlying genetics, biology, clinical characteristics and cellular origins ^1–3^. The most common subtype is high-grade serous OC (HGSOC), which accounts for around two-thirds of all invasive OCs, and likely arises from fallopian tube secretory epithelial cells ^4^. Other subtypes of invasive OC are rarer and include clear cell and endometrioid OCs (CCOC and EnOC), which are strongly associated with the benign precursor lesion endometriosis ^5^, and mucinous OC (MOC), which may derive from appendiceal tissues or primordial germ cells ^6^.

Molecular analyses of primary OCs have so far focused on characterizing somatic genomic variations in protein-coding genes, and have implicated distinct genetic alterations and biological pathways in the development of each histotype. In HGSOCs, *TP53* mutations are ubiquitous ^7^; loss-of-function of DNA double-strand break repair genes (e.g. *BRCA1, BRCA2*) are also common ^8^ and predicate a genomic instability phenotype that results in an accumulation of gross structural genomic changes as tumors develop ^9^. CCOCs and EnOCs often harbor somatic pathogenic mutations in *ARID1A*, a member of the SWI/SNF family of chromatin remodelers ^10^; *TERT* promoter mutations are specific to CCOCs ^11^; and coding mutations and promoter methylation in DNA mismatch repair genes are relatively common in the EnOC subtype ^12^. For MOC, alterations in the *MAPK* pathway are present in∼70% of tumors, with KRAS hotspot mutations (amino acid 12 or 13) the most frequent genetic change ^13^.

There is currently a lack of understanding of the role of non-coding mutations in cancer pathogenesis. Whole genome sequencing (WGS) studies shows that ∼96% of all somatic mutations identified in primary tumors lie in non-protein coding DNA regions. A proportion of non-coding somatic mutations likely represent functional drivers of cancer disease development ^14^, mediating their effects by modifying the function of regulatory elements (REs) that modify the expression of target genes that contribute to neoplastic development. The regulatory architecture of the non-coding genome is highly tissue and cell-type specific ^15^. Somatic mutations within disease-specific REs are therefore expected to affect gene expression in a disease specific manner. The goals of the current study are: (1) To characterize the histotype-specific regulatory landscapes of the different OC histotypes; and (2) using WGS data from primary ovarian tumors, to identify frequently mutated REs (FMREs) that may play a role in disease pathogenesis.

## RESULTS

### Histotype-Specific Epigenomic and Transcriptomic Landscapes of Ovarian Cancers

We characterized the histotype-specific landscapes of active chromatin in primary ovarian tumors using chromatin immunoprecipitation-sequencing (ChIP-seq) for acetylated lysine 27 of histone H3 protein (H3K27ac). H3K27ac ChIP-seq was performed in twenty primary tumors, five each for the different histotypes: HGSOC, CCOC, EnOC and MOC (**Supplementary Table S1**). Using the ENCODE ChIP-seq processing pipeline and quality control guidelines (**Supplementary Figure S1**), we identified a union set of 295,243 non-overlapping ChIP-seq peaks across all tumors, encompassing 11.6% of the genome (**Figure 1a**). The number of new peaks identified saturated at 16 tumors, suggesting we identified the majority of active REs in OC. The 12,954 peaks (4.4%) shared across all tumor types were enriched for promoter regions (odds ratio = 14, P < 0.001, Fisher’s exact test, when compared genome-wide) (**Figure 1b**). For each tumor type, we also identified a set of histotype-specific H3K27ac peaks: 6,583 in HGSOCs, 5,401 in CCOCs, 2,134 in MOCs and 20 in EnOCs. These peaks fall predominantly in enhancers (odds ratio = 40, P<0.001, Fisher’s exact test, histotype-specific H3K27ac peaks *versus* common H3K27ac peaks in OCs) (**Figure 1b**).

**Figure 1.**
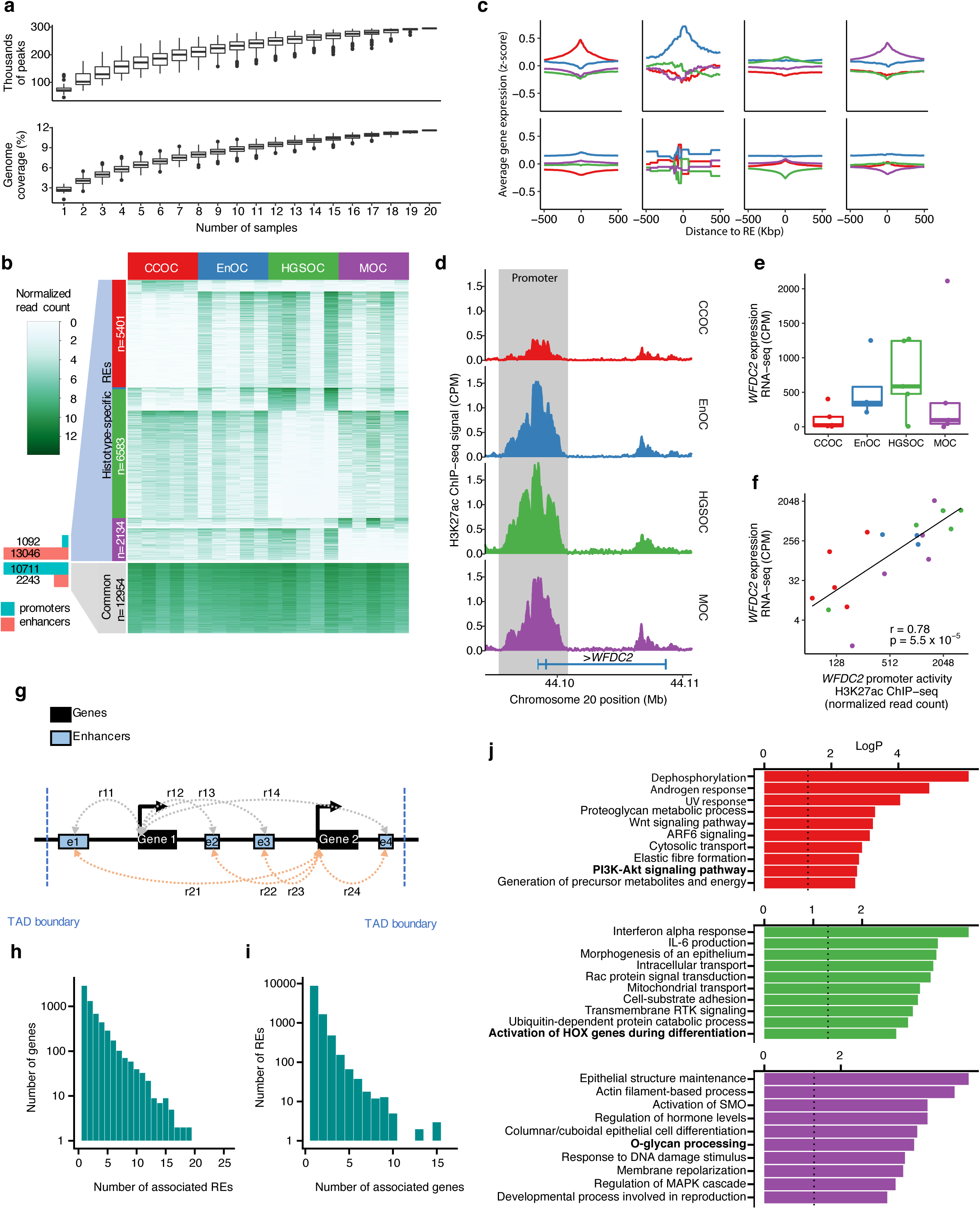
Epigenomic profiling in 20 epithelial OCs reveals histotype-specific REs. (**a**) Number of peaks and genome coverage as a function of number of samples shows 20 OC samples converged in ∼300 thousand peaks that cover 11.6% of the human genome. (**b**) Heatmap showing the normalized H3K27ac ChIP-seq signal (white - absence of H3K27ac peak, light green - weak H3K27ac peak, dark green - strong H3K27ac ChIP-seq signal) for the 20 OC samples (columns) at the active REs (rows). (**c**) Average gene expression around histotype-specific REs showing gene expression is higher in the corresponding histotype in genes around histotype-specific regions that have a positive fold change and vice versa. (**d**-**f**) *WFDC2* locus showing higher H3K27ac ChIP-seq signal in the promoter region (chr20:44,095,981-44,101,008) (**d**) and gene expression (**e**) in EnOC and HGSOC compared to CCOC and MOC. (**f**) Normalized H3K27ac ChIP-seq signal in the promoter of *WFDC2* has a strong correlation with *WFDC2* gene expression. (**g-j**) Enhancer-gene associations using enhancer activity and gene expression contribute to the analysis of histotype-specific enhancers. (**g**) Diagram of the enhancer-gene association strategy that computes the Spearman’s correlation between enhancer activity (normalized H3K27ac ChIP-seq signal) and gene expression (normalized RNA-seq) (r_ij_) between all enhancers and all genes within the same topological associating domain (TAD). A putative enhancer-gene association is established if the correlation (r_ij_) is significant (r_ij_>0.4 and p-value<0.05). (**h**) Histogram of number of associated genes per RE and (**i**) number of associated REs per gene. (**j**) Pathway enrichment analysis of genes associated with histotype-specific REs.

We performed RNA sequencing (RNA-seq) in nineteen of these tumors. Consistent with H3K27ac ChIP-seq data, we identified histotype-specific patterns of gene expression for each tumor type. There were 1,214 differentially expressed genes (DEGs) specific to CCOC, 519 DEGs for HGSOC, 371 DEGs for MOC, and 16 DEGs for EnOC (**Supplementary Figure S2**). Genes flanking histotype-specific peaks of active chromatin were consistently expressed at higher levels in tumors of the same histotype. Conversely, lower H3K27ac signal was associated with lower expression of nearby genes (**Figure 1c**). Genes flanking REs that are shared across the different histotypes show similar patterns of gene expression across the different histotypes (**Supplementary Figure S3**). As an example, we observed elevated H3K27ac ChIP-seq signal in the promoter of *WFDC2* associated with higher expression of the *WFDC2* gene in HGSOC and EnOC samples compared to CCOCs and MOCs (Spearman’s rho = 0.78, P=5.5×10^-5^) (**Figure 1d-f**). *WFDC2* encodes for Human Epididymis Protein 4 (HE4), a biomarker overexpressed in serous and endometrioid OCs ^16^, and is used clinically to monitor OC recurrence. Taken together, these data indicate that histotype-specific enhancers regulate gene expression *in cis* in a histotype-type specific manner.

### Predicting Regulatory Element-Gene Interactions

We integrated H3K27ac ChIP-seq and RNA-seq data to map REs (REs), including typical enhancers and promoters, to putative target genes. We calculated all correlations between gene expression and RE activity to identify all gene-RE pairs within topologically associating domains (TAD) (Spearman’s rho > 0.4, P<0.05, distance < 500Kbp) (**Figure 1g**) ^17^. We use the term ‘CREAG’ (collection of ‘*<u>c</u>is*-<u>R</u>egulatory <u>E</u>lements <u>A</u>ssociated with a <u>G</u>ene’), to define the collection of predicted enhancers for a gene of interest; a CREAG is conceptually similar to gene ‘plexi’ ^18^. This defined a catalogue of 15,380 RE-gene associations between 6,197 genes and 11,371 REs across all histotypes, with a median of 2.5 enhancers per CREAG (**Figure 1h**) and 1.4 genes per enhancer (**Figure 1i**). We compared enhancer-gene associations with the GeneHancer database to infer target genes for 285,000 human enhancers ^19^. Fourteen percent of our predicted enhancer-gene associations were annotated to the same gene in GeneHancer, 2.3 times more than expected by chance (P<0.001). We then mapped all histotype-specific REs to their putative target gene(s) (**Supplementary Table S2**) and performed pathway enrichment analysis to identify biological mechanisms associated with each histotype (**Figure 1j and Supplementary Figure S4**). Genes associated with histotype-specific REs function in pathways known to be involved the development of each histotype. Thus, RE-associated genes for CCOC (e.g. *AKT2, ITGA5, LAMC1, MET, PPP2R3A* and *SGK1*) are involved in PI3K-Akt signaling (P=0.0017) ^3,20,21^; and active REs specific to MOCs are associated with genes (e.g. *GCNT3, MUC12, GALNT5* and *B3GNT5*) involved in O-glycan processing (P=0.0001), which may reflect upregulated mucin production and *O*-glycosylation activity in this histotype ^22^. Pathways common across histotypes were associated with cell proliferation and mitosis (**Supplementary Figure S4**).

### Super-enhancer Landscapes in Ovarian Cancers

Large enhancer domains, termed super-enhancers (SEs) or stretch enhancers ^23^, typically regulate genes critical to cell identity. We asked whether histotype-specific SEs were present, and whether these were associated with genes that may contribute to phenotypic heterogeneity in OCs. We characterized between 653 and 1,945 SEs per tumor (mean = 1,245, sd = 321) (**Figure 2a**, **Supplementary Table S3**). By assigning SEs to the highest expressed overlapping gene we identified a total of 5,338 SE-associated genes (SEAGs), of which 1,123 were common to all OC histotypes. *PAX8,* a transcription factor and known essential gene in OC ^24^, was associated with SEs in 17 (out of 20) tumors, with the lowest signal in MOCs. *MUC16*, which encodes CA125, another clinical biomarker used to monitor OC, coincided with an SE in 13 (out of 20) tumors; again the lowest enhancer activity was in MOCs, which are known to have lowest of CA125 expression levels compared to other histotypes ^25^ (**Figure 2b and Supplementary Figure S5a**). Novel SEAGs for OC include the highly expressed lncRNA *LINC00963,* which was identified in 17 (out of 20) tumors (average expression = 470 CPM, sd = 585 CPM, **Supplementary Figure S5b**). *LINC00963* has been implicated in prostate cancer progression ^26^.

**Figure 2.**
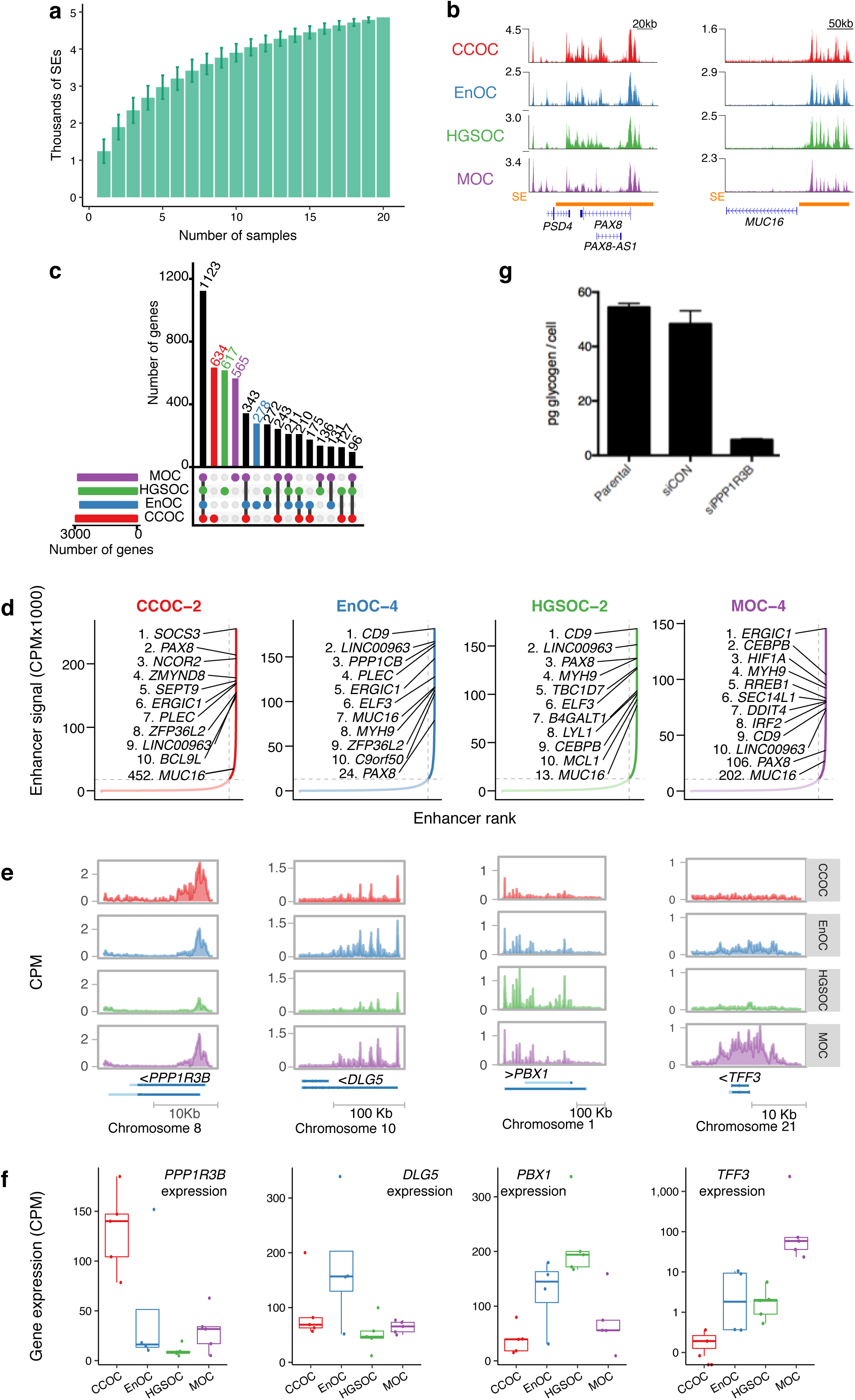
Histotype-specific super-enhancers. (**a**) Number of super-enhancers as a function of number of samples. (**b**) *PAX8* and *MUC16* loci show common OC SEs. (**c**) UpsetR plot showing the size of the all the subsets of SEAGs by whether they are present in CCOC, EnOC, HGSOC and MOC. (**d**) Gene expression of the histotype-specific SEAGs show higher expression levels for the histotype of interest. (**e**) Plots of all enhancers ranked by enhancer signal for four representative OC samples, one for each histotype, showing the top 10 SEAGs, and the rank of *PAX8* SE, *MUC16* SE and *EPCAM* SE for each sample. (**f-g**) Examples of histotype-specific SEAGs. Gene tracks (**f**) and gene expression (**g**) that show one example of histotype-specific SEAGs for each histotype (*PPP1R3B, DLG5, PBX1*, and *TFF3* for CCOC, EnOC, HGSOC and MOC). (**h**) Glycogen absorbance in JHOC5 before and after PPP1R3B knockdown.

A total of 2,094 SEAGs were histotype-specific: 634 for CCOC, 617 for HGSOC, 565 for MOC and 278 for EnOC (**Figure 2c**). As expected, histotype-specific SEAGs were more highly expressed in the histotype of interest (**Supplementary Figure S5c**); for example, SEAGs specific to CCOC had a median normalized gene expression above zero, but less than zero for all other histotypes. The known function of many of the SEAGs we identified support a histotype-specific role in OC (**Figure 2d-f**); for example the CCOC specific gene Protein Phosphatase 1 Regulatory Subunit 3B (*PPP1R3B*) regulates glycogen synthesis, consistent with the observations that clear cell tumors contain large deposits of cytoplasmic glycogen. *In vitro PPP1R3B* knockdown in a CCOC model (JHOC5 cell line) results in a significant decrease in glycogen content (P=0.05, **Figure 2g**). SEAGs common to all OCs are involved in pathways such as TNFA signaling via NF-kB and response to growth factor (P=2.7 x 10^-25^ and 1.3 x 10^-14^, respectively, (**Supplementary Figure S5d**).

### Somatically Mutated Regulatory Elements in Primary Ovarian Cancers

We collated WGS data from 232 OCs (169 HGSOCs, 28 CCOCs and 35 EnOCs) to identify the non-coding somatic single nucleotide variants (SNVs) occurring in active enhancers and promoters in OCs. In total, there were 1.7 million somatic SNVs across all 232 tumors, with an average of 7,163 SNVs per tumor (range 480-40,576) (**Supplementary Figure S6a**). Of these, 1.6 million (98.8%) SNVs lay in non-coding DNA regions (**Supplementary Figure S6b**). We identified all somatic non-coding SNVs that intersect consensus H3K27ac regions in ovarian tumors. Approximately 9.3% percent of SNVs were located in active REs. About 14% of SNVs were in active promoters and 86% were in active enhancers. We identified frequently mutated regulatory elements (FMREs) harboring somatic SNVs at a frequency greater than expected by chance based on the average distribution throughout the genome (see Methods). There were 25 FMREs across all histotypes at a false discovery rate (FDR) of 0.25 (**Figure 3a-c**): 8 FMREs were unique to HGSOC and 17 were unique to endometriosis-associated OCs (CCOC and EnOC) (**Supplementary Table S4**). FMREs included promoters and enhancers. Using our gene-RE maps, we quantified global patterns of differential gene expression associated with somatic SNVs in promoters and enhancers. Based on a subset of 89 HGSOCs for which both WGS and RNA-seq data were available, we found overall that genes were significantly more likely to be overexpressed in samples with RE mutations compared to wild-type samples, i.e., samples without somatic mutations in the RE of interest. For the 2,893 REs harboring at least 1 somatic SNV, 89 genes were significantly overexpressed (FC > 2), and 46 genes were significantly under expressed (FC < 0.5) (p-value<0.05, binomial distribution), indicating enhancer SNVs are more likely to activate rather than repress gene expression (**Figure 3d**). On chromosome 10, we identified a cluster of nine somatic SNVs all located within the *KLF6* promoter, part of a common SE in primary OCs (**Figure 3e**). Two variants coincided with a binding site for PAX8 in the promoter defined by H3K27ac-ChIP-seq in OC cell lines ^27^. SE activity is associated with *KLF6* expression in all OC histotypes, but the SE is only mutated in HGSOC (P=8.2 x 10^-8^). *KLF6* is a Krüppel-like transcription factor with tumor suppressor functions, and is associated with chemoresponse and prognosis in OC patients ^28^.

**Figure 3.**
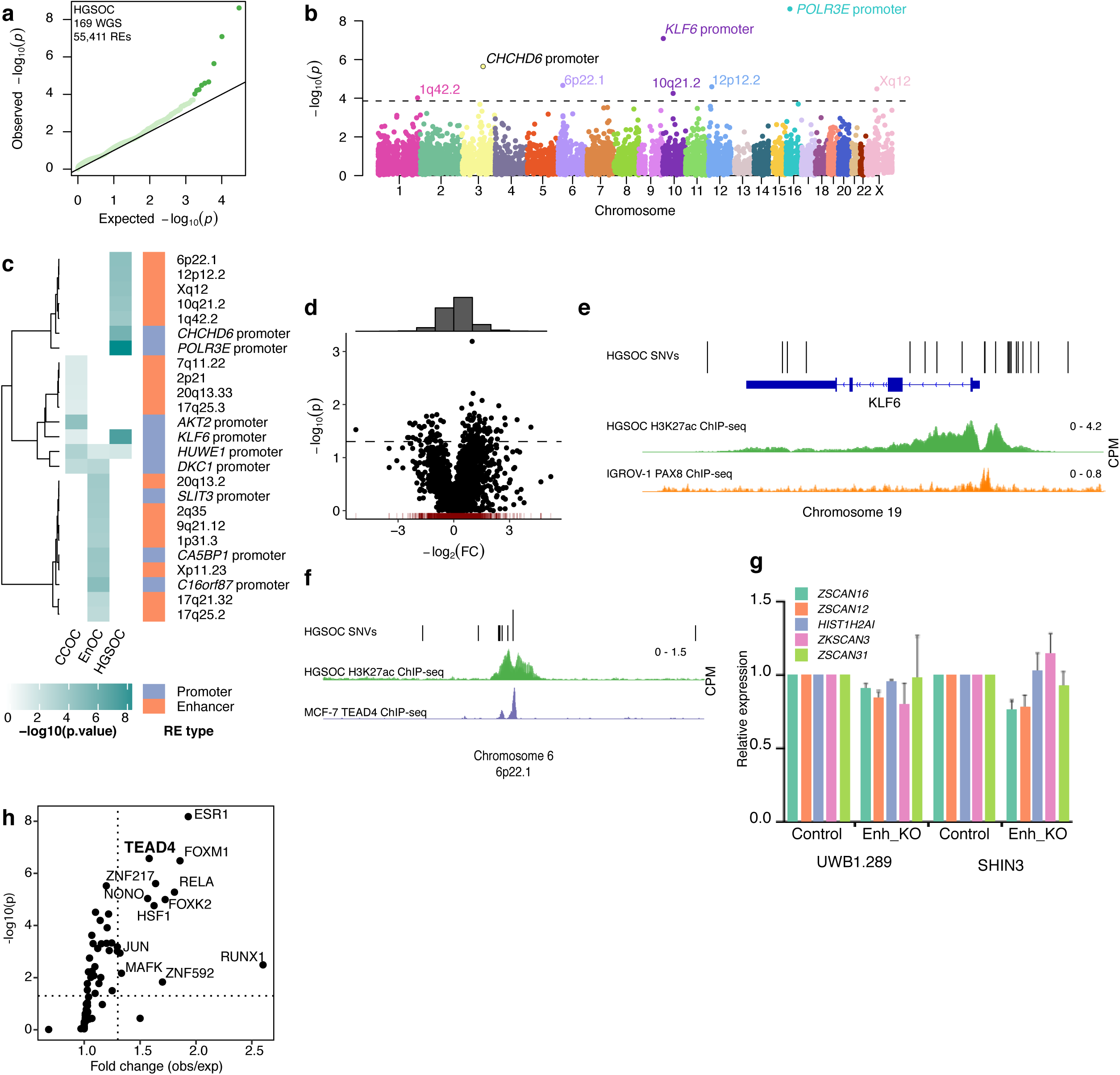
Frequently mutated regulatory elements (FMREs) in OC. (**a**) Heatmap that shows the mutational burden (p-value) of the 25 FMREs across CCOC, EnOC and HGSOC. (**b**) Manhattan plot that shows the genomic location (x-axis) and significance value (y-axis) of the mutational burden for all active REs in HGSOC. (**c**) QQ-plot that shows the expected (x-axis) and observed (y-axis) significance values of the mutational burden for all active REs in HGSOC. (**d**) Volcano plot that shows the fold change of median gene expression (x-axis) and the significance value (y-axis) of the putative target gene of samples with overlapping single nucleotide variants in an active RE vs. wild-type samples. The histogram on top of the scatterplot shows more overexpression events (-log2(FC) > 0) in the presence of single nucleotide variants than under expression events (-log2(FC) < 0). (**e**) The KLF6 promoter is frequently mutated in high-grade serous OC. Mutations cluster closer to the promoter region of KLF6 and overlap PAX8 BSs in OC cell lines. (**f**) An enhancer in the 6p22.1 locus is frequently mutated in high-grade serous OC. The normalized H3K27ac ChIP-seq signal of the enhancer shows the strongest correlation (Spearman’s rho = 0.69, P=6.6 x 10^-4^) with *ZSCAN16* gene expression, which is more than 200 Kb away, and null correlation to nearby genes. The single nucleotide variants within the enhancer fall in valleys of the H3K27ac ChIP-seq peak and overlap TEAD4 BSs. (**g**) Relative expression of *ZSCAN16, ZSCAN12, HIST1H2AI, ZKSCAN3* and *ZSCAN31* before and after the knockout of the FMRE enhancer at 6p22.1 locus. *ZSCAN16* expression is statistically significant decreased after enhancer knockout, validating our *in silico* prediction. **(h)** TEAD4 BSs are frequently mutated in high grade serous OC. Diagram showing the fold change (observed/expected number of samples with at least one mutation in the BSs overlapping the consensus set of active enhancers and promoters) vs. the p-value (significance of the observed number of mutated samples) showing TEAD4 and other TFs have more mutations than expected by chance in the set of active BSs,.i.e., BSs that are within H3K27ac ChIP-seq peaks.

At the 6p22.1 locus, we identified a cluster of seven somatic SNVs in an enhancer located ∼9kb centromeric to the *HIST1* gene cluster (**Figure 3f**). In our dataset, the activity of this putative enhancer correlates strongly with the expression of *ZSCAN16* (Spearman’s rho = 0.69, P=6.6 x 10^-4^) and *ZSCAN12* (Spearman’s rho = 0.47, P=0.02). We used CRISPR/Cas9 to knock out ∼630bp of this enhancer in two HGSOC cell lines (UWB1.289 and SHIN3). Enhancer knockout induced downregulation of both *ZSCAN16* and *ZSCAN12* (P=0.05) (**Figure 3g**), while the expression of three other genes within the TAD were not significantly affected, including *ZKSCAN3* and *HIST1H2A1*, additional putative targets of this enhancer ^29^. *ZSCAN16* or *ZSCAN12* are TFs containing SCAN domains that mediate protein-protein interactions; neither have been previously implicated in OC and very little is known about their function.

Using motifBreakR ^30^ we predicted that a somatic SNV (chr6:27870735:T>A) in 6p22.1 enhancer identified in two different tumors breaks a TEAD4 motif. Using TEAD4 ChIP-seq data generated in MCF-7 breast cancer cells, we found TEAD4 binding sites overlap 4/8 (50%) FMREs identified in HGSOCs, including the 6p22.1 enhancer. Active TEAD4 binding sites (coinciding with H3K27ac ChIP-seq OC peaks) were significantly mutated in ovarian tumors (fold change(obs/exp) = 1.8, P=6 x 10^-11^) (**Figure 3h**). Globally, we observed that mutations in TEAD4 binding sites occur preferentially in the active TEAD4 binding sites – of 10,872 and 1,968 TEAD4 binding sites and active binding sites 1.8% and 9.4%, respectively, harbored a somatic SNVs in ovarian tumors. TEAD4 is a known binding partner of PAX8 ^31,32^. Consistent with this we found enrichment of somatic SNVs in HGSOCs in active PAX8 binding sites (FC = 1.9, P=2×10^-10^). One hundred and nine of 169 HGSOCs (64.5%) harbored a somatic SNV in at least one active PAX8 binding site. Taken together these data indicate that somatic point mutations converge on TEAD4/PAX8 binding sites to deregulate PAX8 target gene expression during OC progression.

### Gene-centric Analysis of Frequently Mutated Regulatory Elements

We tested the aggregate of somatic mutations in 6,197 CREAGs identified in primary ovarian tumors. Nine hundred and sixteen genes had a significantly increased mutation burden across multiple REs associated for each gene (P<0.05, **Figure 4a-c**). There was statistically significant enrichment of SE associated genes among the most frequently mutated CREAGs (P=0.01, **Figure 4d**) which included genes known to be involved in OC pathogenesis; for example hepatocyte nuclear factor 4, gamma (*HNF4G*), homeobox D9 (*HOXD9*) ^33^, N-myc downstream regulated 1 (*NDRG1*) ^34^, CD47 molecule (*CD47*) ^35^, MDS1 and EVI1 complex locus (*MECOM*) ^36^ and *PAX8* ^27^.

**Figure 4.**
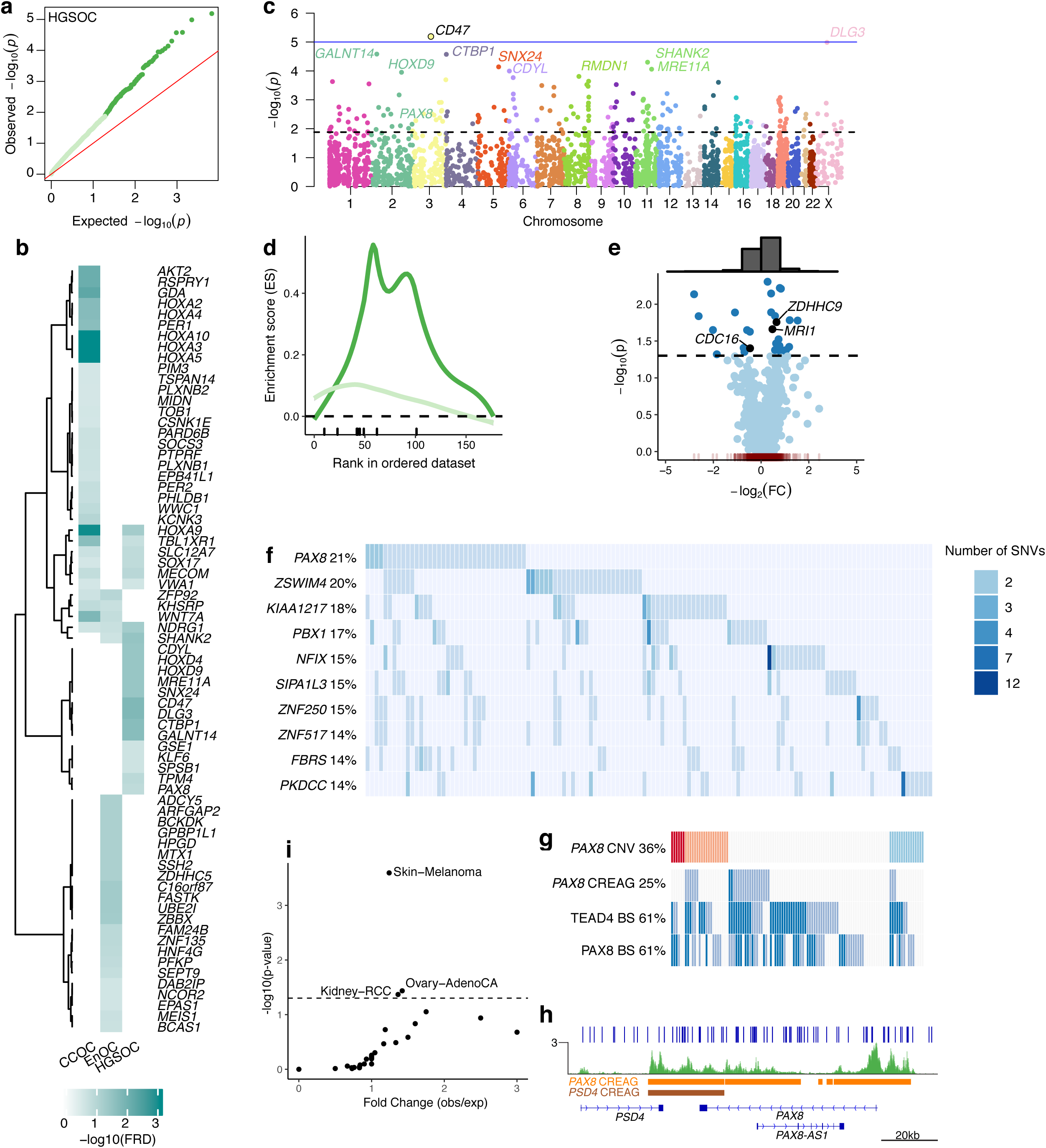
Gene-centric mutation rate aggregated across the collection of REs associated to a gene (CREAG). (**a**) QQ-plot that shows the expected (x-axis) and observed (y-axis) significance values of mutational burden for all CREAGs. (**b**) Super-enhancer associated genes that have somatic SNVs. (**c**) Manhattan plot showing the genomic location (x-axis) and significance (y-axis) of mutational burden for all CREAGs. (**d**) Enrichment of SEAGs in FM CREAGs (dark green) vs. random selection (light green) (P=0.01). (**e**) Volcano plot that shows the fold change of median gene expression (x-axis) and the significance value (y-axis) of the putative target gene of samples with overlapping single nucleotide variants in a CREAG vs. wild-type samples. The histogram on top of the scatterplot shows more overexpression events (-log2(FC) > 0) in the presence of single nucleotide variants than under expression events (-log2(FC) < 0). (**f-h**) PAX8 CREAG is frequently mutated in OC. (**f**) HGSOC (n = 169) non-coding oncoplot showing the top 10 ranking genes in terms of number of samples with mutations overlapping its CREAG showing PAX8 as the highest mutated CREAG. (g) HGSOC (n=110) oncoplot with PAX8 copy number variation, PAX8 CREAG mutation, TEAD4 and PAX8 BS mutations. (h) PAX8 locus showing the mutations on PAX8 CREAG. (**i**) The number of samples with mutations overlapping the PAX8 CREAG are statistically significant in Skin-Melanoma, Ovary-Adeno-CA and Kidney-RCC datasets.

We measured differential gene expression as a function of mutational state for all associated REs for each gene. Comparing between samples with mutated CREAGs and samples with wild-type CREAGs, we found thirty significantly differentially expressed genes (Mann Whitney U test P<0.05). More of these genes were associated with gain of function (n = 20) compared to loss of function (n = 10). Overexpressed genes include prothymosin-alpha (*PTMA)* (FC = 1.26, P=0.005, Mann Whitney U test); integrator complex subunit 1 (*INTS1*) (FC = 1.44, P=0.01, Mann Whitney U test); and NOC2 like nucleolar associated transcriptional repressor (*NOC2L*) (FC = 1.66, P=0.01, Mann Whitney U test). Down regulated genes included lysophosphatidic acid receptor 3 (*LPAR3*) (FC = 0.09, P=0.007, Mann Whitney U test) and metallothionein 1X (*MT1X*) (FC = 0.4, P=0.013, Mann Whitney U test) (**Figure 4e**). Analysis of gene essentiality in 517 cancer cell lines (including 53 OC cell lines) show that *PTMA, INTS1* and *NOC2L* are essential genes in cancer ^37^.

Based on the normalized mutational burden (p-value) and the frequency with which somatic SNVs occur in any of the REs that contribute to a CREAG, the *CD47* CREAG (containing 5 REs) is the most significantly associated with OC and contains 22 mutations in 21 patients (P=6.46×10^-6^, **Figure 4c**). Overall, the *PAX8* CREAG was the most frequently mutated in HGSOC; 36/169 tumors (21%) harbored a somatic mutation in at least one of three REs in this CREAG (P=0.003) (**Figures 4f-h**). PAX8 CREAG is also mutated in other cancer types, however, it is only statistically significant in skin cancer, kidney cancer and ovarian cancer (**Figure 4i**).

## DISCUSSION

Different OC histotypes can be defined by distinct protein-coding somatic driver events and have different disease etiologies; but little is known about the underlying mechanisms of gene regulation and their contribution to disease pathogenesis for the different histotypes and their precursor tissues. To our knowledge, this study describes for the first time, the histotype-specific regulatory architecture of OC based on H3K27ac ChIP-seq analysis of primary tumors representing the four major subtypes of invasive disease. As anticipated, different histotypes of OCs share common epigenomic features consistent with their common site of disease development; but we show conclusively that each subtype also has an RE signature that delineates the transcriptional mechanisms that underlie the development of each histotype. Endometrioid OCs (EnOCs) represent an exception in that H3K27ac ChIP-seq analysis only identified a small number of histotype-specific REs. When we focused on the histotype-specific REs, 2 of our EnOCs resembled HGSOC, while the other 3 resembled CCOC. This is consistent with clinical and biological traits of these tumors; EnOCs are more biologically related to clear cell OCs (CCOCs); but late-stage EnOCs can be histopathologically misdiagnosed as high-grade serous ovarian OCs (HGSOCs). This combination of shared etiology and possible misdiagnosis may be explanation for the lack of specificity in defining the regulatory architecture this histotype.

Histotype-specific REs were strongly enriched for putative enhancers. Enhancer depletion was more common than enhancer gain, consistent with previous reports that loss of activity drives cell-type specific identity. A more in-depth analysis of enhancer activity identified around 1,100 ‘Müllerian’ SEs that are common across all OC histotypes. SEs mark genes associated with cell lineage and cell state, in normal and cancer tissues alike ^38^. We identified SEs coinciding with genes that encode established biomarkers in OC, highlighting the disease-specific nature of these findings. This included SEs associated with mucin 16 (*MUC16*) that encodes CA125, and *PAX8*. *MUC16*/CA125 is a reliable serum marker that is used clinically to aid in the diagnosis of OC and monitor disease progression. *PAX8* is a lineage specific TF that is highly expressed in fallopian tube epithelia, a precursor of HGSOC, is amplified in ∼80% of HGSOCs, and is an essential gene in OC ^8,24^. The *PAX8* SE was detected in all OC histotypes and most active in HGSOC and EnOC. A SE proximal to *PPP1R3B* was unique to CCOC, *PPP1R3B* was highest expressed in CCOC compared to the other histotypes, and *PPP1R3B* knockdown in a CCOC model confirmed its role in production of the glycogen-rich cytoplasm that characterizes CCOC. Similarly, we found other histotype-specific SEs which likely regulate genes important for establishing the defining features of each OC histotype. Overall, the data indicate a role for SE landscapes as underlying drivers of ovarian and histotype-specific pathogenesis.

By integrating mutation data from whole genome sequencing (WGS) of ovarian tumors into these analyses we were able to identify somatic mutations that fall into REs to identify candidate noncoding REs that may contribute to disease pathogenesis. Most ovarian tumors contain several thousand somatic mutations, the vast majority of which (98.7%) lie in the non-protein coding DNA regions; but only a proportion of these are likely to have a functional impact on disease development. Because there is no genetic code for the non-coding genome, one needs to address several possible hypotheses in these analyses: (1) that somatic mutations perturb specific REs, including active enhancers and super-enhancers, to affect *in cis* the expression of gene(s) that are critical in disease biology; (2) that the constellation of somatic mutations in multiple REs within TADs affect local gene expression; and (3) that the burden of somatic mutations within lineage/disease specific REs (e.g. PAX8 binding sites) contribute to affect the expression within transcriptional networks.

Even though these studies are currently restricted by the numbers of tumors for which data are available from WGS analysis (232 invasive ovarian tumors), a summary of our findings strongly indicate that somatic non-coding mutations in REs play an important role in disease etiology. Firstly, frequently mutated regulatory elements (FMREs) showed histotype-specificity, consistent with the histotype-specificity observed from integrated ChIP-seq and RNA-seq analysis. Second, FMREs we associated with genes that are known to be biologically important in OC development. Finally, burdens of somatic mutations in REs were predicted to deregulate pathways that are critical in OC biology. Inevitably, these studies will benefit in the future from additional WGS analyses performed in ovarian tumors and in greater numbers for the different histotypes. There are also likely to be limitations in the current analyses because we understandably focused on somatic SNVs and assumed that all SNVs have a similar functional impact on an RE. Because of restrictions in statistical power we were not able to evaluate other somatic variation that may contribute to disrupting gene regulation, including aneuploidy, large interstitial deletions and rearrangements, and microdeletions/amplifications, all which vary widely between tumors and could affect one to several hundred REs.

Perhaps the most compelling finding was the convergence of several analyses that underscore the importance of PAX8 regulation in OC development, in particular for the high-grade serous subtype. We observed a significant clustering of mutations in PAX8-associated upstream enhancers in HGSOCs but in CCOC or EnOC. PAX8-bound enhancers were also significantly mutated in HGSOC, as were enhancers bound by TEAD4, a known PAX8 binding partner ^27,31^, indicating a role for PAX8 as both a target of and mediator of non-coding somatic mutations. We predict that functional somatic mutations in REs alter binding of a sequence-specific DNA-binding protein, such as a TF, which was suggested by the finding of somatic mutations disrupting the TEAD4 motif within a mutated enhancer on chromosome 6. Knockout of the frequently mutated enhancer at 6p22.1 locus, validated a positive association between the activity of the enhancer and *ZSCAN16* expression. Further analysis is needed to determine whether there is a direct contact between the RE and *ZSCAN16*.

Our studies also identified novel genes and transcription factors in OC pathogenesis that warrant additional functional studies to validate their role in cancer. In addition to PAX8- and TEAD4-bound enhancers, we also identified significantly increased mutation rates in FOXM1 and ESR1 bound regions in HGSOC. Analysis of HGSOC TCGA data indicates FOXM1 is a frequently altered pathway in HGSOC development ^8^ and ESR1 is expressed in around 80% of HGSOCs. The somatic mutations at these transcription factor binding sites, could affect the binding specificity forming either stronger or weaker binding, altering their downstream pathways.

In conclusion, through the integration of tissue-specific epigenomic and gene expression landscapes for the different histotypes of OC, and somatic mutation data from large scale whole genome sequencing analyses of primary tumors, we have identified non-coding elements that may contribute to OC development. Many of the mutated REs and their associated genes provide novel insights into disease pathogenesis, and the lineage-specific TF PAX8 was identified a central player in the transcriptional dysregulation caused by non-coding somatic SNVs in HGSOCs.

## METHODS

For more detailed procedures please refer to the **Supplementary Methods**.

### Tissue ChIP-Seq

All tissues used were collected with informed consent and the approval of the institutional review boards of University of Southern California and Cedars-Sinai Medical Center. Tissue ChIP-Seq was performed as previously described ^39^. Chromatin was immunoprecipitated using an anti-H3K27ac antibody (DiAGenode, C15410196, Denville, NJ). Sequencing libraries prepared from enriched DNA using the ThruPLEX-FD Prep Kit (Rubicon Genomics, Ann Arbor, MI) and were sequenced using 75-base pair single reads on the Illumina platform (Illumina, San Diego, CA) at the Dana-Farber Cancer Institute. The AQUAS pipeline ^40^ was used to processed all H3K27ac ChIP-Seq data, with histotype-specific regions called using Diffbind ^41^.

### RNA-Seq

Primary ovarian cancer specimens were homogenized and total RNA was extracted using TRIzol LS (Thermo Fisher Scientific, catalogue number: 10296028). Ribosomal RNA (rRNA) was depleted using RiboMinus Transcriptome Isolation Kit (Thermo Fischer Scientific, catalogue number: K155002). Poly (A)+ RNA was then isolated using Dynabeads Oligo (dT) 25 (Thermo Fischer Scientific, catalogue number: 61002). Twenty nanograms rRNA-poly (A)+ RNA was used to prepare each RNA-Seq library. External RNA Controls Consortium (ERCC) spike-ins (Thermo Fischer Scientific, catalogue number: 4456740) were added as control for normalization of the samples. Strand-specific RNA-Seq libraries were constructed using the NEBNext Ultra Directional RNA Library Prep Kit (NEB, catalogue number: E7420). The resulting library concentrations were quantified using the Nanodrop. Libraries were sequenced to generate paired-end 75 bp reads on NextSeq 500 platform (Illumina) in high output running mode. Sequencing was performed at the Molecular Genomics Core facility at the University of Southern California.

### RE/gene mapping

We calculated the pairwise correlation (*r*) of the H3K27ac ChIP-Seq score of a given RE against the gene expression, in counts per million (CPM), of all genes within the same TAD. For the remaining RE/gene pairs (Bonferroni-adjusted p < 0.05), we computed a transformed correlation value (*r*′_*ij*_) as a function of the distance between the *i* th RE and *j* th gene, as follows:

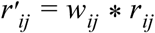

where *r*_*ij*_ is the correlation between the *i* th RE and *j* th gene, and *w*_*ij*_ is a weight parameter calculated as follows:

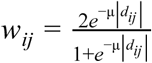

where *d*_*ij*_ is the distance between the *i* th RE and the *j* th gene (in base pairs) and μ is a parameter set such as *w*_*ij*_ equals 0.5 when *d*_*ij*_ is 200 kbp. We then discarded all RE/gene associations with an *r*′_*ij*_ > 0.5, restricting automatically all RE/gene associations to be within 200 kbp, and including distal associations if the original correlations (*r*_*ij*_) are close to 1, but “weaker” correlations if the RE is close to the gene.

### Identifying Frequently Mutated Regulatory Elements

We used SNVs from 232 OCs. To identify FMREs we used a Poisson binomial distribution with a vector of probabilities, i.e., one background mutation rate for each sample. When *X*_*i*_ (*X*_*i*_∈[0, 169]) is a random variable that represents the number of samples with at least one mutation in the *i* th RE, then *X*_*i*_ follows a Poisson binomial distribution (PBD) with a vector of probabilities 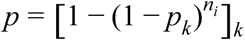, where *n*_*i*_ is the size of the *i* th RE in base pairs, and *p*_*k*_ is the global background rate of sample *k* (*k*∈[1, 169]) empirically estimated by the ratio of the total number of non-coding SNVs in sample *k* (*n*_*k*_) over the total coverage of the regions of interest in base pairs, (the H3K27ac-positive regions) (*n*_*cov*_):

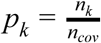

To determine whether the observed number of mutated samples in the *i* th RE (*s*_*i*_), we calculate the probability of having at least *s*_*i*_ samples mutated, i.e., p-value _i_ = *P* (*X*_*i*_ ≥ *s*_*i*_). P-values were adjusted used the Benjamini-Hochberg method. For gene-level analysis, we combined all REs and annotated promoters associated to gene *j* into a pseudo RE and estimated the probability of having *s*_*j*_ or more mutated samples in all combined REs, i.e., p-value _j_ = *P* (*X*_*j*_ ≥ *s*_*j*_), where *X*_*j*_ follows a PBD with a vector of probabilities 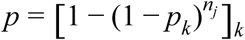, where *n*_*j*_ is the total length of the pseudo RE, i.e., the sum of the widths of all REs and annotated promoters associated to gene *j*.

### Quantifying Gene Expression Changes Associated with RE Mutations

For 89 WGS samples in PCAWG, RNA-Seq was available. Analysis was restricted to genes with at least two mutated samples in the RE overlapping the promoter or an associated RE. For each gene, we calculated the fold change (FC), defined as the ratio of the median gene expression of the gene in the mutated samples (x_1_) over the median gene expression of the gene in the non-mutated samples (x_2_). Pathway enrichment analysis was performed using Metascape, using an adjusted p-value cutoff of 0.10.

## ACKNOWLEDGEMENTS

This work was supported by a Tower Cancer Research Foundation Career Development Award and a K99/R00 grant from the National Cancer Institute (NCI) (1K99CA184415-01) to K.L. The tissue specimens were either collected as part of the USC Jean Richardson Gynecologic Tissue and Fluid Repository, which is supported by a grant from the USC Department of Obstetrics & Gynecology and the NCT Cancer Center Shared Grant award P30 CA014089 (to the Norris Comprehensive Cancer Center) or as part of the Women’s Cancer Biobank at Cedars-Sinai Medical Center. This work was supported in part by the Ovarian Cancer Research Fund Alliance Program Project Development Grant (373356): Co-Evolution of Epithelial Ovarian Cancer and Tumor Stroma (B.Y.K, K.L.).

