## Supplementary Figures for "Non-coding Somatic Mutations Converge on the PAX8 Pathway in Epithelial Ovarian Cancer"

### Slide 1
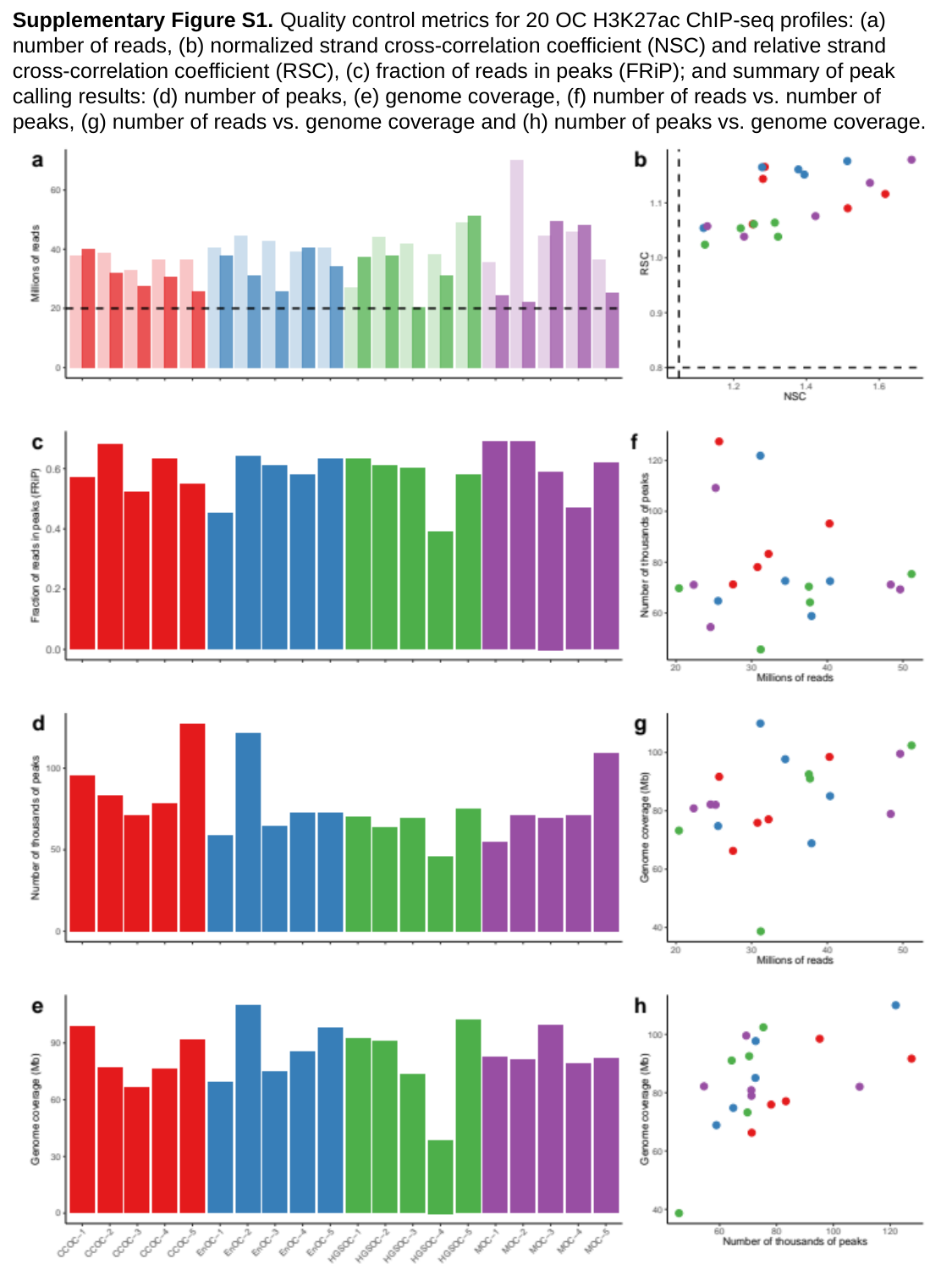

Supplementary Figure S1. Quality control metrics for 20 OC H3K27ac ChIP-seq profiles: (a) number of reads, (b) normalized strand cross-correlation coefficient (NSC) and relative strand cross-correlation coefficient (RSC), (c) fraction of reads in peaks (FRiP); and summary of peak calling results: (d) number of peaks, (e) genome coverage, (f) number of reads vs. number of peaks, (g) number of reads vs. genome coverage and (h) number of peaks vs. genome coverage.

### Slide 2
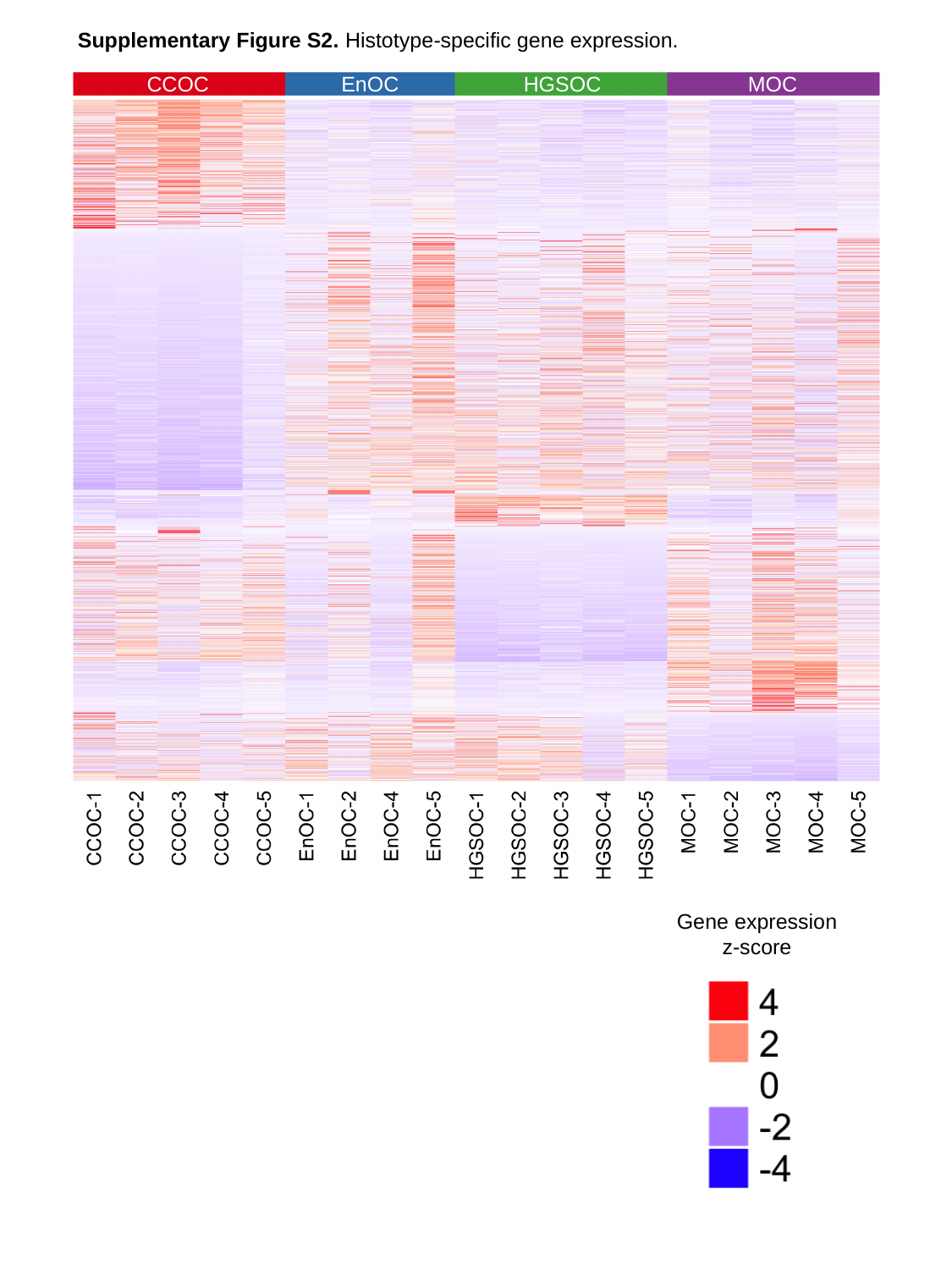

Supplementary Figure S2. Histotype-specific gene expression.
CCOC
EnOC
HGSOC
MOC
Gene expression
z-score

### Slide 3
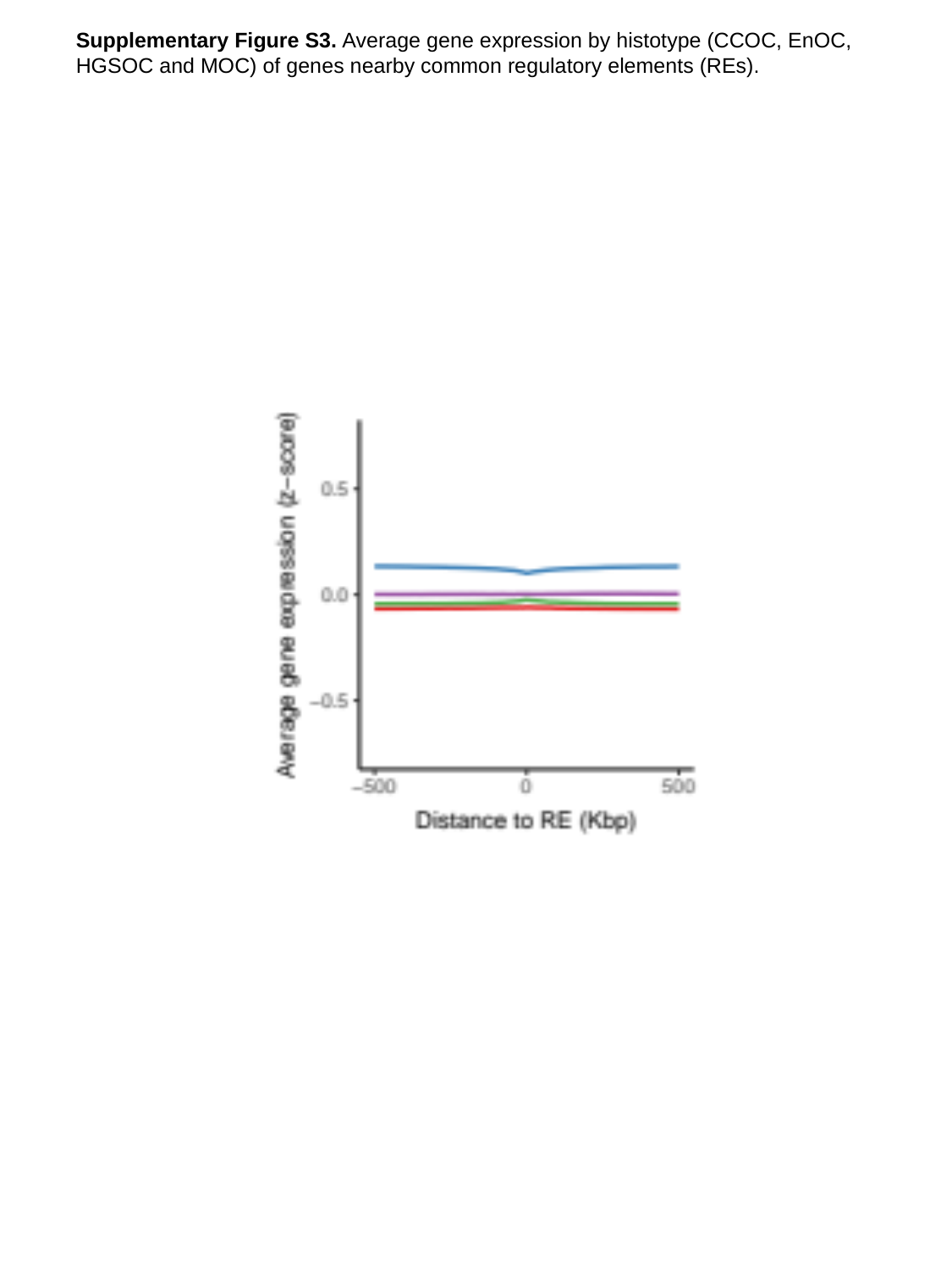

Supplementary Figure S3. Average gene expression by histotype (CCOC, EnOC, HGSOC and MOC) of genes nearby common regulatory elements (REs).

### Slide 4
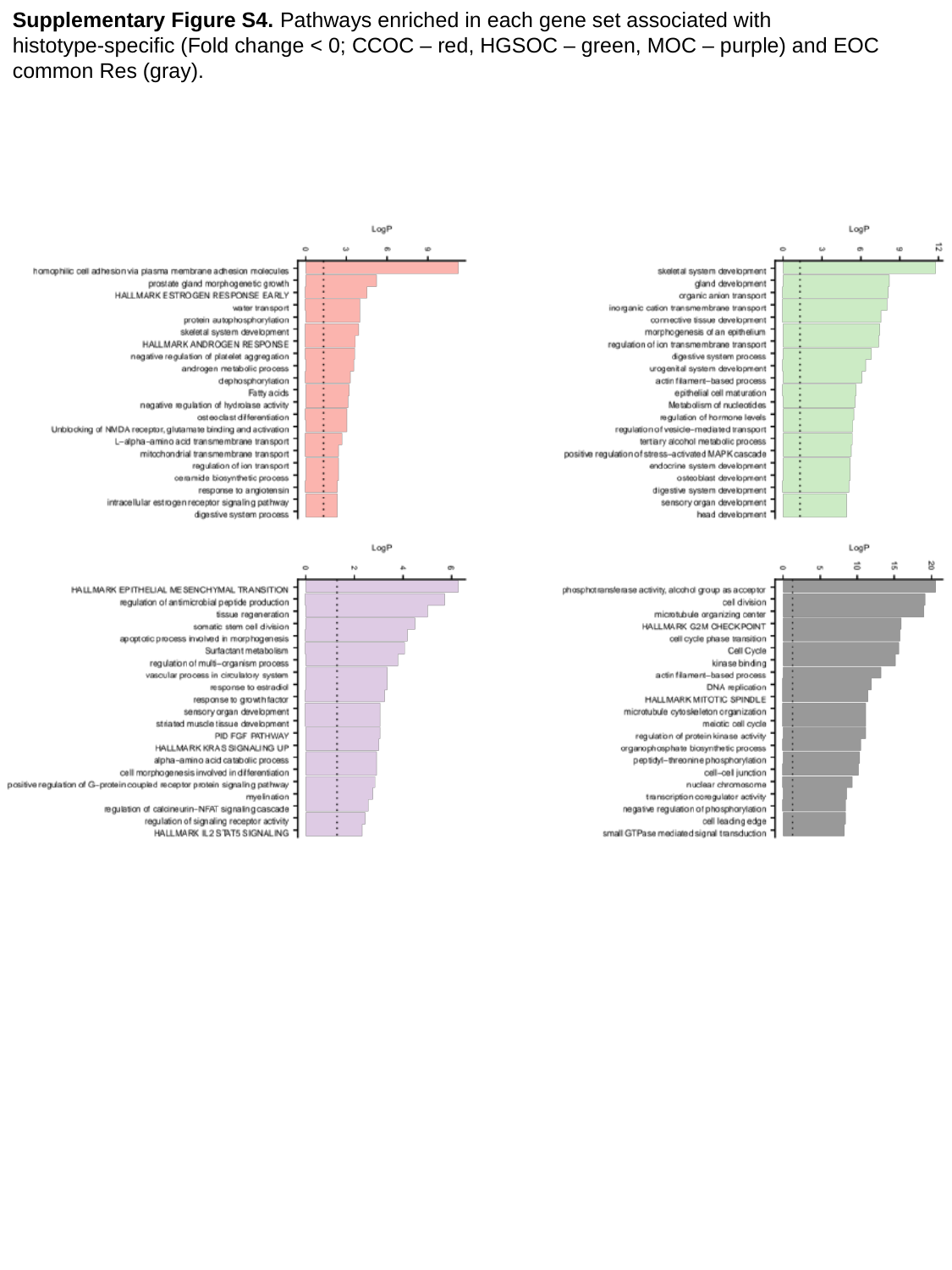

Supplementary Figure S4. Pathways enriched in each gene set associated with
histotype-specific (Fold change < 0; CCOC – red, HGSOC – green, MOC – purple) and EOC common Res (gray).

### Slide 5
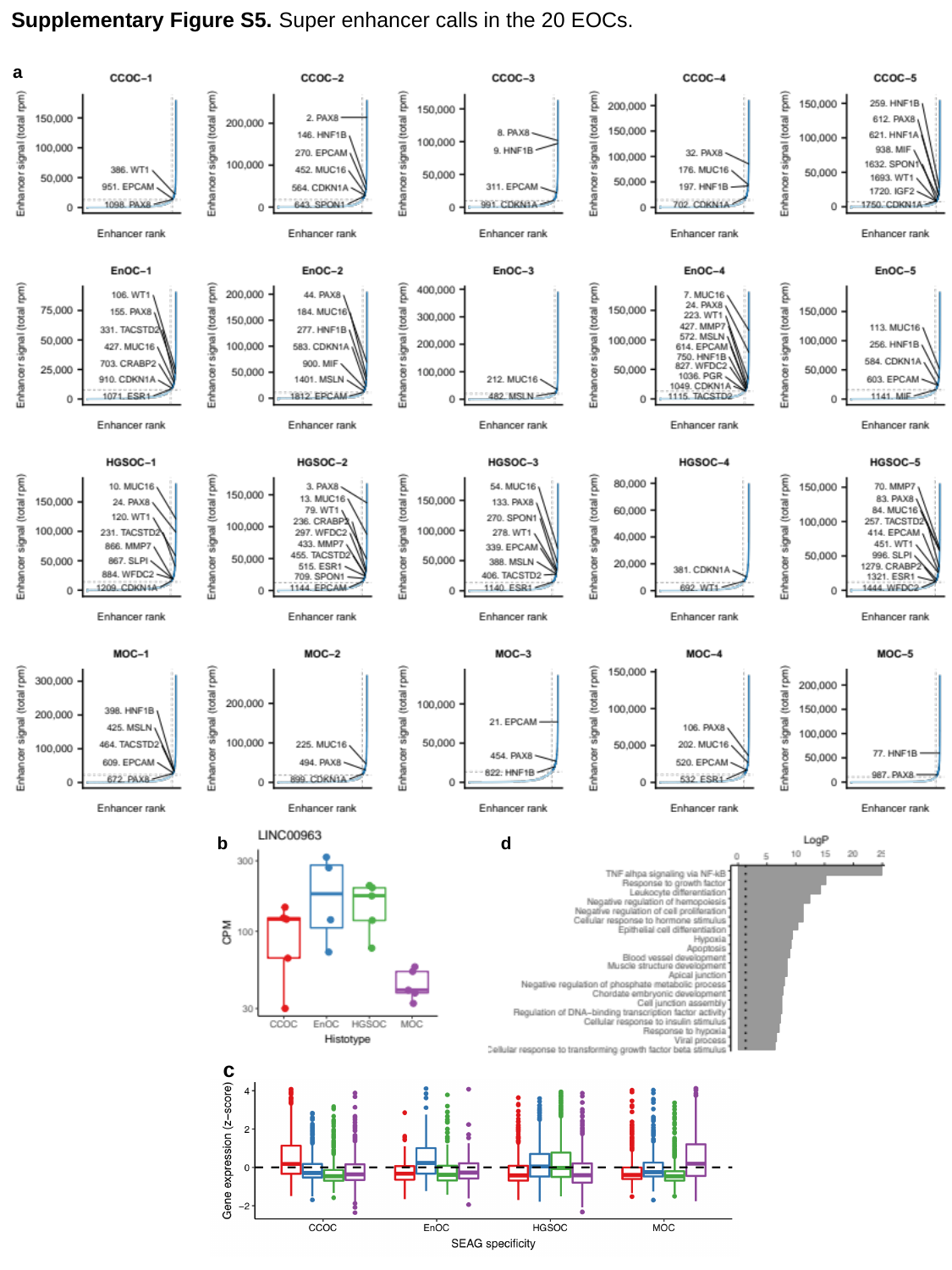

Supplementary Figure S5. Super enhancer calls in the 20 EOCs.
a
d
b
c

### Slide 6
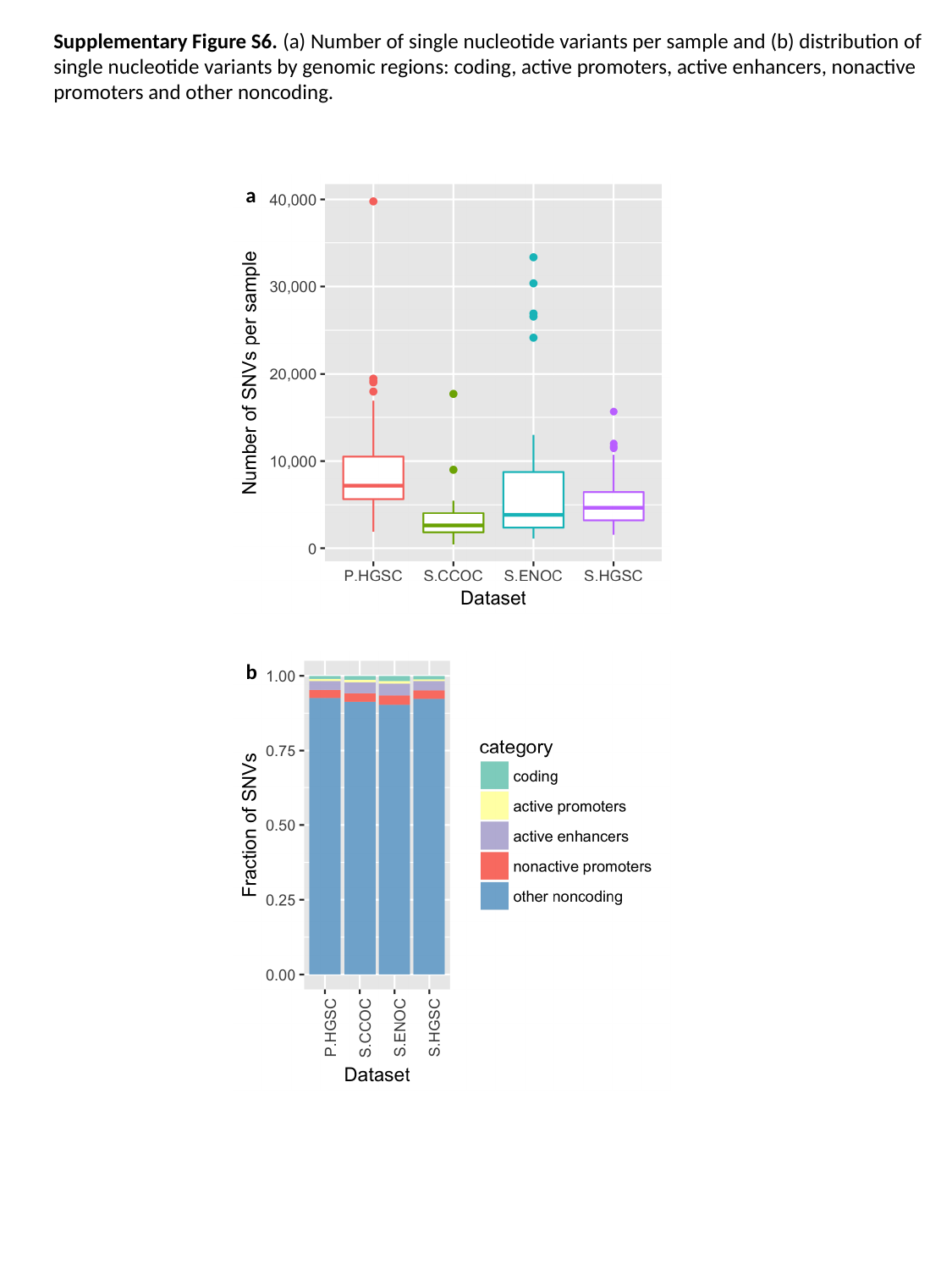

Supplementary Figure S6. (a) Number of single nucleotide variants per sample and (b) distribution of
single nucleotide variants by genomic regions: coding, active promoters, active enhancers, nonactive
promoters and other noncoding.
a
b
