## Supplementary Methods for "Non-coding Somatic Mutations Converge on the PAX8 Pathway in Epithelial Ovarian Cancer"

Corona *et al.,*

**SUPPLEMENTARY METHODS**

***Tissue ChIP-Seq***

All tissues used were collected with informed consent and the approval of the institutional review boards of University of Southern California and Cedars-Sinai Medical Center. Tissue ChIP-Seq was performed as previously described [(32)](http://f1000.com/work/citation?ids=1638024&pre=&suf=&sa=0). Briefly, one frozen 3 mm core was pulverized in a pulverization bag (TT05) using the Covaris CryoPrep system (Covaris, Woburn, MA) twice at intensity 4. The tissue was then fixed using 1% formaldehyde (Thermo fisher, Waltham, MA) in Phosphate-buffered saline solution for 10 minutes at room temperature with rotation, and quenched with 125 mM glycine for 10 min at room temperature with rotation. After rinsing with ice-cold Phosphate-buffered saline solution twice, chromatin was resuspended and lysed in ice cold lysis buffer (50 mM Tris, 10 mM EDTA, 1% SDS with protease inhibitor) for 10 minutes. Chromatin was sheared to 300-500 base pairs using the Covaris E210 sonicator (AFA: 5% duty cycle, 5 intensity, 200 cycles/burst) for 10 min. 1% of chromatin was saved as input for each sample. 5 vol of dilution buffer (1% Triton X-100, 2 mM EDTA, 150 mM NaCl, 20 mM Tris HCl pH 8.1) was added to the rest of chromatin and the sample was incubated with 1 µg H3K27ac antibody (DiAGenode, C15410196, Denville, NJ) coupled with protein A and protein G beads (Life Technologies, Carlsbad, CA) at 4°C overnight. The chromatin was washed with RIPA washing buffer (0.05M HEPES pH 7.6, 1 mM EDTA, 0.7% Na Deoxycholate, 1% NP-40, 0.5M LiCl) for 5 times and, rinsed with TE buffer (pH 8.0) once. The sample was resuspended in elution buffer (50 mM Tris, 10 mM EDTA, 1% SDS), treated with RNase for 30 minutes at 37°C, and incubated with proteinase K overnight at 65°C. Sample DNA and input were extracted using Qiagen Qiaquick columns, and sequencing libraries prepared using the ThruPLEX-FD Prep Kit (Rubicon Genomics, Ann Arbor, MI). Libraries were sequenced using 75-base pair single reads on the Illumina platform (Illumina, San Diego, CA) at the Dana-Farber Cancer Institute.

**ChIP-seq data processing**

Reads were aligned against the reference human genome hg19, filtered by quality and duplication. Several quality control metrics were computed for each individual replicate, including number of reads, percentage of duplicated reads, NSC, RSC and FRiP (Supplementary Figure S1). AQUAS follows ENCODE3 guidelines to process ChIP-Seq data. For all cell line ChIP-Seq experiments, we had two biological replicates. For histone modification ChIP-Seq data, AQUAS uses macs2 as the peak caller algorithm and a naive overlap approach that selects for the final peakset the regions of the pooled replicate that overlaps 50% or more of each individual replicate. <describe number of peaks of individual replicates and overlap of individual replicates>

For the 20 OCs, we obtained an average of 33.7 million mapped reads (standard deviation, sd = 9,071,589), and an average of 77,346 peaks (sd = 21,020), per sample. H3K27ac peaks were, on average, 1078 bp wide (sd = 1206 bp), and covered around 83 Mbp per sample (sd = 16 Mbp) (Supplementary Figure S1).
There was negative correlation between the number of peaks and the average peak width (Pearson’s ⍴ = -0.58, P-value = 0.007) and positive correlation between number of peaks and genome coverage (Pearson’s ⍴ = 0.54, P-value = 0.007).

Histotype-specific regions were identified using the R bioconductor package DiffBind [(34)](http://f1000.com/work/citation?ids=181771&pre=&suf=&sa=0), which calculates differentially bound regions from multiple ChIP-seq experiments. For each histotype, we selected sites called in at least 3 out of 5 samples in the given histotype, with absolute Fold change value greater than or equal to 3, FDR less than or equal to 0.05, contrasting the 5 samples of the histotype of interest against the remaining 15 samples using the method DBA_EDGER.

Common regions that were called in all 20 OC H3K27ac ChIP-Seq experiments regardless of the intensity of the ChIP-Seq signal. The consensus set of H3K27ac ChIP-seq regions for HGSOC were all sites present in at least 3 (out of 5) HGSOC samples, merging overlapping peaks and we recalculated the ChIP-Seq score using DiffBind with the new coordinates across all samples, to have homogenous start/end positions for all peaks across the samples.

**RNA-seq data analysis**

Reads were aligned to hg38 (ref_genome_hg38_gencodev26) using STAR (STAR-2.5.1b), then a read count matrix was generated using featureCounts (gencode.v24.annotation.gtf). The samples range from 8.6 to 32.2 million of uniquely mapped reads.

Histotype-specifically expressed genes were identified using the R Bioconductor package DESeq2 with absolute log_2_ FC value greater or equal to 2, p-value less or equal to 0.05, contrasting the 5 samples (4 in case of EnOC) of one histotype against the remaining 14 (15 in case of EnOC).

**Filtering single nucleotide variants**

We used single nucleotide variants (SNVs) from 169 HGSOCs (110 from the PCAWG cohort and 59 from the UBC cohort). We removed all SNVs that overlap coding regions, regions annotated as low mappability regions by ENCODE (wgEncodeDacMapabilityConsensusExcludable.bed), as well as areas of the genome that are prone to cause false positives in ChIP-Seq assays (seq.cov1.ONHG19.bed.gz). For HGSOC, there is a total of 1’276,929 SNVs that fall into 1’276,108 individual positions (1275309 positions with only 1 mutated sample, 784 positions with two mutated samples, 11 positions with three mutated samples, 3 positions with four mutated samples, and one position (chr3:174499208-174499208) with 7 mutated samples).

**Generation enhancer-deletion mutant in ovarian cancer cell lines**

CRISPR/Cas9 system was used to generate deletion mutant. FUCas9Cherry (Gifts from Dr. Marco Herold, Addgene plasmid number 70182) was transfected together with lentiviral packaging plasmids pMD2.G and psPAX2 (Gifts from Dr. Didier Trono, Addgene plasmid numbers 12259 & 12260) into 293T cells. UWB1.289 and SHIN3 cells were then transduced with lentivirus and mCherry-positive cells (UWB1.289/Cas9 and SHIN3/Cas9) were sorted by FACS. Cas9 activity was confirmed using gRNAs targeting the RB1 locus, and Surveyor T7 Endonuclease 1 digests. UWB1.289/Cas9 and SHIN3/Cas9 were subsequently transduced by lentivirus containing either gRNA pair targeting enhancer region on chromosome 6 (Chr6-3) or gRNA pair targeting a control gene OR1C1 before selection with 400 ng/mL puromycin. Plasmids expressing gRNAs were obtained from Transomic Technology. Sequences for Chr6-3 gRNA-A are: TCCCTTGCCAGCTCACTCAA; Chr6-3 gRNA-B: AGGAATCCAACTAATACCAT; OR1C1 gRNA-A: AGGGCTGAAATAGACGGCGA; OR1C1 gRNA-B: AGAGGTGATCTTCAGAACAG.
